# Hypoxia/HIF Signaling Negatively Regulates Bone Marrow Adiposity after Radiation Exposure

**DOI:** 10.1101/2025.09.07.674016

**Authors:** Cheyenne A. Jones, Wendi Guo, Kiana A. Gunn, Cahil Potnis, Amaya Sheffield, Colleen Wu

## Abstract

Radiation therapy is an essential cancer treatment, yet collateral damage to normal tissues remains a major clinical challenge. In bone, radiation-induced toxicity is characterized by loss of hematopoietic function, reduced bone volume, and increased marrow adipose tissue (MAT). Importantly, cancer patients who undergo radiotherapy exhibit significantly higher fracture risk compared to those receiving similar treatments without radiation exposure, underscoring the clinical consequences of bone microenvironment (BME) injury. The BME is inherently hypoxic resulting in the activation of by hypoxia-inducible factor (HIF) signaling. Here, we demonstrate that radiation induces a rapid and persistent accumulation of MAT, with adipocytes localizing preferentially to hypoxic regions of the marrow. To investigate the role of hypoxia/HIF signaling in this process, we generated *aP2Cre;Hif-1 ^fl/fl^;Hif-2^fl/fl^* conditional knockout mice. Surprisingly, these mice exhibited increased MAT expansion following radiation compared to controls, suggesting that HIF deletion in aP2-expressing cells exacerbates radiation-induced adipogenesis. Analysis aP2CreRosa26^tdTomato/+^ mice revealed that most aP2-expressing cells did not give rise to mature adipocytes, macrophages, or endothelial cells, pointing instead to an uncharacterized stromal population that influences MAT formation. In contrast, conditional ablation of HIFα transcription factors in *LepRCre*-expressing skeletal stem cells, which contain a subpopulation of skeletal progenitors which directly contribute to marrow adipocytes, had no effect on radiation-induced MAT expansion. Collectively, these findings identify a previously unrecognized population of adipocyte-regulatory cells whose HIF-dependent activity constrains stress-induced marrow adiposity. This work provides new mechanistic insight into how radiation disrupts the marrow microenvironment and expands MAT, advancing our understanding of the cellular and molecular drivers of radiation-induced bone fragility.

**One Sentence Summary:** Loss of HIF signaling in aP2Cre expressing cells enhances radiation induced marrow adiposity.

## INTRODUCTION

Bone loss associated with radiation therapy is attributed, in part, to a transient increase in osteoclast activity and suppression of osteoblast actively (1, 2, 3, 4, 5). Unfortunately, strategies focused on targeting cells of the bone remodeling unit to prevent osteopenia have not significantly decreased risk for fractures, nor improved hematopoietic function for cancer patients (6, 7, 8). Importantly, expansion of marrow adipose tissue (MAT) is associated with diminished hematopoietic stem cell (HSC) frequency and decreased bone volume (9, 10, 11). For these reasons, identifying novel mechanisms to that contribute to radiation induced MAT may aid in preventing insufficiency fractures and improve hematopoiesis in patients undergoing radiotherapy

Pharmacologic activation of the HIF signaling pathway mitigates gastrointestinal tissue toxicity, in part, by preventing vascular damage through the secretion of VEGF (12). However, HIF signaling is highly context dependent with cell types responding uniquely to activation (13, 14). While the BME contains regional areas of hypoxia, the role of HIF signaling in regulating MAT has not been characterized (15, 16, 17, 18). Notably, in obesity models, genetic ablation of HIF transcription factors in white adipose tissue (WAT) prevents adipocyte hypertrophy (19). Collectively, this supports the concept that modulation of the HIF signaling pathway can be used to ameliorate MAT formation after radiation exposure to bone. Recent advances in microscopy (16) and the development of oxygen sensitive probes (15) for live imaging support the concept that the BME contains regional areas of hypoxia (17, 18). Under hypoxic conditions, HIF-1α and HIF-2α transcription factors are stabilized and coordinate the cellular response to hypoxia by activating gene expression programs that facilitate oxygen delivery and cellular adaptation to oxygen deprivation (20)

Targeted deletion of HIFα in cells within the osteoblast lineage demonstrate HIF signaling couples angiogenesis to osteogenesis by regulating VEGF expression to control bone formation (21, 22). Furthermore, *in vivo* loss of function and gain of function show that HIF signaling in osteoblast cells regulates osteoblast differentiation in a HIF-1α dependent manner and bone remodeling via expression of OPG in HIF-2a dependent manner (14, 23). HIF signaling also regulates the spatio-temporal onset of angiogenesis in the growth plate where HIF-1 plays a central role in hypoxia-dependent cartilage formation and maintenance (24), whereas HIF-2α is involved in endochondral ossification and cartilage destruction [53]. Thus, HIF signaling is active in osteoblasts and chondrocytes with isoform specific functions of HIF-1α and HIF-2α for promoting bone formation and cartilage development. While the BME contains regional areas of hypoxia, the role of HIF signaling in regulating MAT has not been characterized (15, 16, 17, 18). Notably, in obesity models, genetic ablation of HIF transcription factors in white adipose tissue (WAT) prevents adipocyte hypertrophy (19). Collectively, this supports the concept that modulation of the HIF signaling pathway can be used to ameliorate MAT formation after radiation exposure to bone.

Multipotent mesenchymal progenitor cells (MMPs) residing in the bone microenvironment (BME) directly contribute to marrow adipocytes (25, 26). Although these studies reveal the cellular origins of MAT, cells that indirectly support adipogenesis in the BME have not been characterized. Recent data from single cell RNA sequencing studies of WAT revealed a subpopulation of adipocyte regulator cells within the vascular stroma fraction which can suppress adipocyte formation in a paracrine fashion (27). Intriguingly, many secreted factors that regulate adipogenesis are putative HIF target genes (28, 29, 30, 31, 32, 33). Given the highly heterogenous and unstructured nature of the bone marrow stroma, the characterization of cells which support adipocytes in the BME represents a gap in knowledge that needs to be filled.

## RESULTS

### Rapid expansion of marrow adipocytes after radiation exposure in a dose and time dependent manner

To define the response of marrow adiposity to radiation exposure, 12-16 week old B6 mice were exposed to either 2, 3, 4 or 6 Gy of total body irradiation (TBI). 7 days post radiation exposure, histological analysis of the distal femur revealed a dose dependent increase in marrow adipocyte number, marrow adipose area and marrow adipose area/marrow area (**FIG 1 A-C**). As we noted a statistically significant increase in adipocyte number beginning after 3 Gy TBI when compared to non-irradiated controls, we performed histological examination and/or microCT quantification of osmium tetra oxide staining and qPCR analysis of long bones 1,3,7, and 14 days post irradiation Consistent with our previous trial, statistically significant differences in MAT were not observed until 7 days post irradiation (**FIG 1 D&E**). In addition, increases in mRNA expression of *Cebpa* and *Pparg,* genes which are classically associated with adipogenesis peaked 3 post IR (**FIG 1 F&G**). Importantly, as alterations in weight can influence MAT content, we did not note significant differences in weight between irradiated and control mice (**FIG S1A**). In addition to MAT expansion, IR is also associated with decreased bone volume, interestingly, while we noted MAT expansion beginning at day 7, bone loss was not observed until 14 days post IR (**FIG S1B**). These results were also observed in both WT mice exposed to 6 Gy of TBI and 8Gy single limb IR demonstrating nonspecific impacts are not responsible for the observed phenotypes. Collectively, we demonstrate that radiation exposure results in a dose and time dependent increase in MAT.

**Figure 1.**
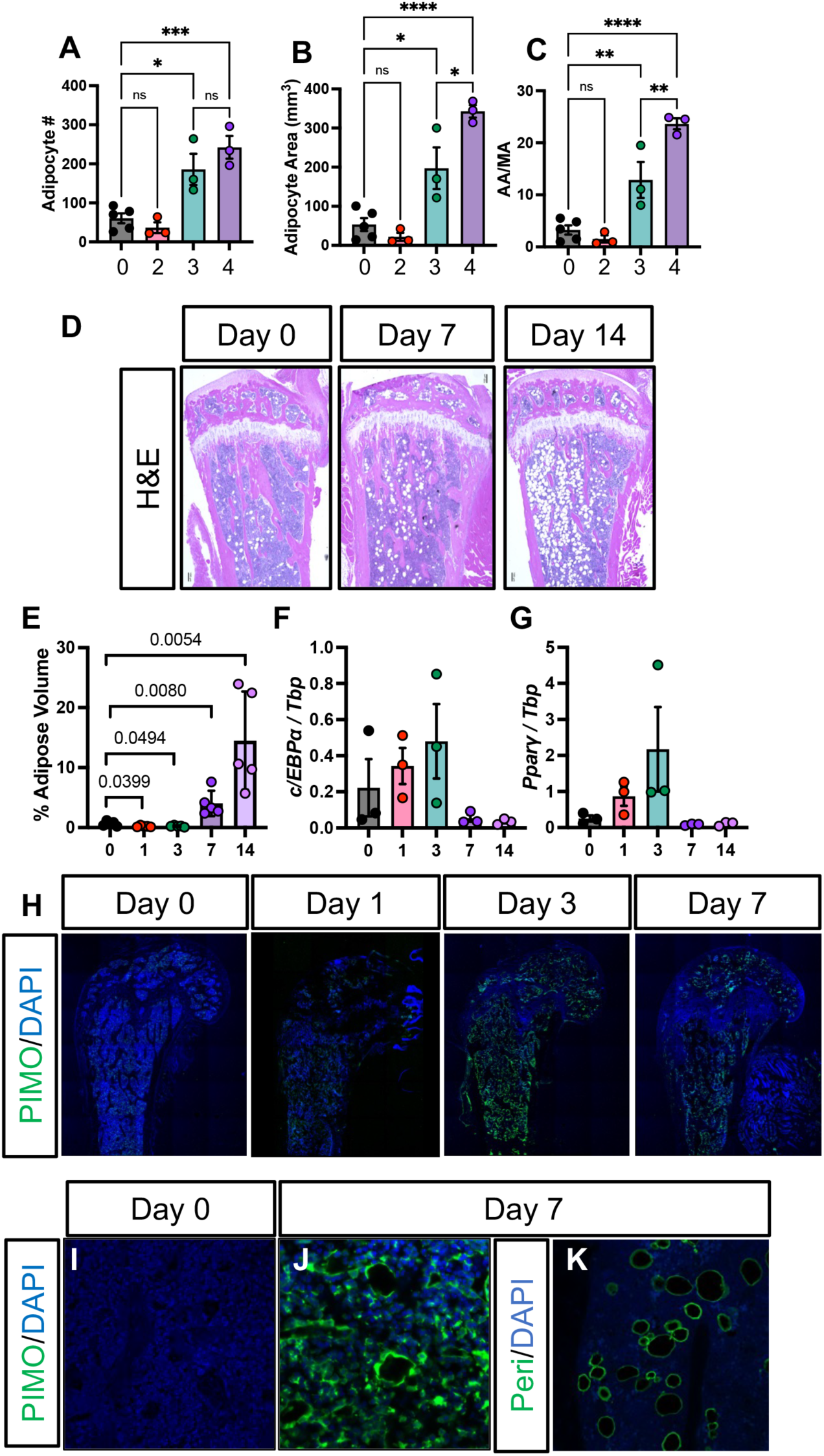
Increased marrow adipose tissue (MAT) after radiation exposure. **(A-C)** Histological analysis of 12-16 week B6 mice after exposure to 2, 3, or 4 Gy of total body irradiation 7 days post (TBI). Quantification of (A) adipocyte number, (B) adipocyte area, and (C) adipocyte number/marrow area. (**D**) Representative H&E images of proximal tibias isolated from 12-16 week old B6 mice 7 and 14 days post IR (**E)** microCT quantification of osmium tetroxide staining of distal femurs and (**F-G**) qPCR analysis of whole bone homogenates after 1, 3, 7, and 14 days post IR. (**H - J**) Representative images of confocal analysis of distal femurs stained for PIMO Days and (**K**) Perilipin. Statistical analysis was performed using one-way ANOVA with Tukey’s post hoc test. Data are presented as means ± SEM. \**p* < 0.05, \*\**p* < 0.01, \*\*\**p* < 0.001, and \*\*\*\**p* < 0.0001.

### Marrow adipocytes are found in hypoxic regions after IR

We have previously demonstrated that radiation exposure disrupts the local endothelium and this vascular swelling is associated with enhances the hypoxic landscape in the BME. However, the contribution of hypoxia/HIF signaling in radiation induced MAT expansion has not been extensively investigated. Towards this end, to examine the characterize MAT in relation to hypoxic status in the BME, we injected mice with pimoniadazole (PIMO) an agent that binds to hypoxic regions to allow for IF visualization of hypoxia. Consistent with previous reports, confocal analysis revealed 1,3, and 7 days post radiation shows that at both 3 and 7 dyas post IR, an increase of PIMO staining when compared to non-irradiated controls (**FIG 1F**). Perilipin staining demonstrated the presences of marrow adipocytes and interestingly, pimonidazole positive staining cells that morphologically resemble marrow adipocytes (**FIG 1G**). This suggests that marrow adipocytes are associated in areas of low oxygen after IR in the BME.

### aP2 Cre expressing cells are expand after IR and are found in hypoxic regions

Previous studied have demonstrated that hypoxia/HIF signaling ameliorates WAT expansion in high fat diets. Based on our observations that after IR adipocytes are found in regions of hypoxia, we sought to determine if HIF signaling was required for marrow adipocyte expansions. The *aP2* gene which encodes for fatty acid binding protein (Fabp4) was originally identified as an adipocyte specific protein (42, 43, 44). When crossed to reporter mouse lines, *aP2Cre* transgenic mice demonstrate adipocyte-specific recombination (45, 46, 47). Expression of the aP2Cre transgene has not been previously characterized in bone marrow adipose tissue which is unique in both function and developmental origins relative to other fat depots. For this reason, we generated *aP2CreRosa26^tdTomato/+^*conditional reporter mice and evaluated the specificity of Cre mediated recombination of *aP2Cre* expressing cells in the hypoxic BME after radiation exposure (**FIG 2A**). Imaging analysis of pimonidazole revealed tdTomato^+^ cells localized to hypoxic regions of the in the bone marrow stroma (**FIG 2B&C**). Confirming these studies, flow cytometry analysis of bone marrow flushes isolated from *aP2CreRosa26^tdTomato/+^* mice 3 days post irradiation tdTomato^+^PIMO^+^ confirmed that the majority of aP2Cre expressing cells reside in regions of hypoxia (**FIG 2D**). Moreover, these studies revealed an expansion of ap2Cre expressing cells in irradiated *aP2CreRosa26^tdTomato/+^* when compared to non-irradiated controls (**FIG 2E&F**).

### Conditional ablation of *Hif-1α* and *Hif 2α* in aP2Cre expressing cells increases MAT after IR

Supporting a role for hypoxia/HIF signaling in stress induced adipose tissue formation, conditional ablation of HIF transcription factors in aP2Cre expressing cells reduced WAT expansion, protecting animals from high fat diet induced obesity(19). For these reasons, we crossed *aP2Cre* mice were crossed to *Hif-1^fl/fl^;Hif-2^fl/fl^*mice to generate *aP2Cre;Hif-1^fl/fl^;Hif-2^fl/fl^* conditional knock out mice. PCR analysis of adipocyte genomic DNA of revealed the presence of recombined (1-Lox) allele indicating efficient activity of CRE recombinase in *aP2Cre;Hif-1^fl/fl^;Hif-2^fl/fl^* cKO mice (**FIG S2A-C**). To determine if HIF is required for MAT expansion in aP2Cre expressing cells, *aP2Cre*(*-*) *and aP2Cre;Hif-1^f/+l^;Hif-2^fl/+^* control and *aP2Cre;Hif-1^fl/fl^;Hif-2^fl/fl^* cKO mice were exposed to 3 Gy TBI. Similar to studies examining WAT, genetic ablation of HIFa proteins in aP2 expressing cells did not impact homeostatic MAT (**FIG 2G,I**) (19). Surprisingly, histological analysis and microCT quantification of long bones isolated from *aP2Cre;Hif-1^fl/+^;Hif-2^fl/+^*/*aP2Cre*(-) control and *aP2Cre;Hif-1^fl/fl^;Hif-2^fl/fl^*cKO animals 7 days post irradiation revealed that conditional ablation of HIF transcription factors increased radiation induced MAT formation in male mice but not female mice (**FIG 2G,I; S2D**). Consistent with our initial studies in WT mice (**FIG 1**), at this early time point, statistically significant differences in bone volume, trabecular separation, trabecular thickness, and trabecular number were not detected in male or female mice (**FIG 2J-L; S2E-G**).

**Figure 2.**
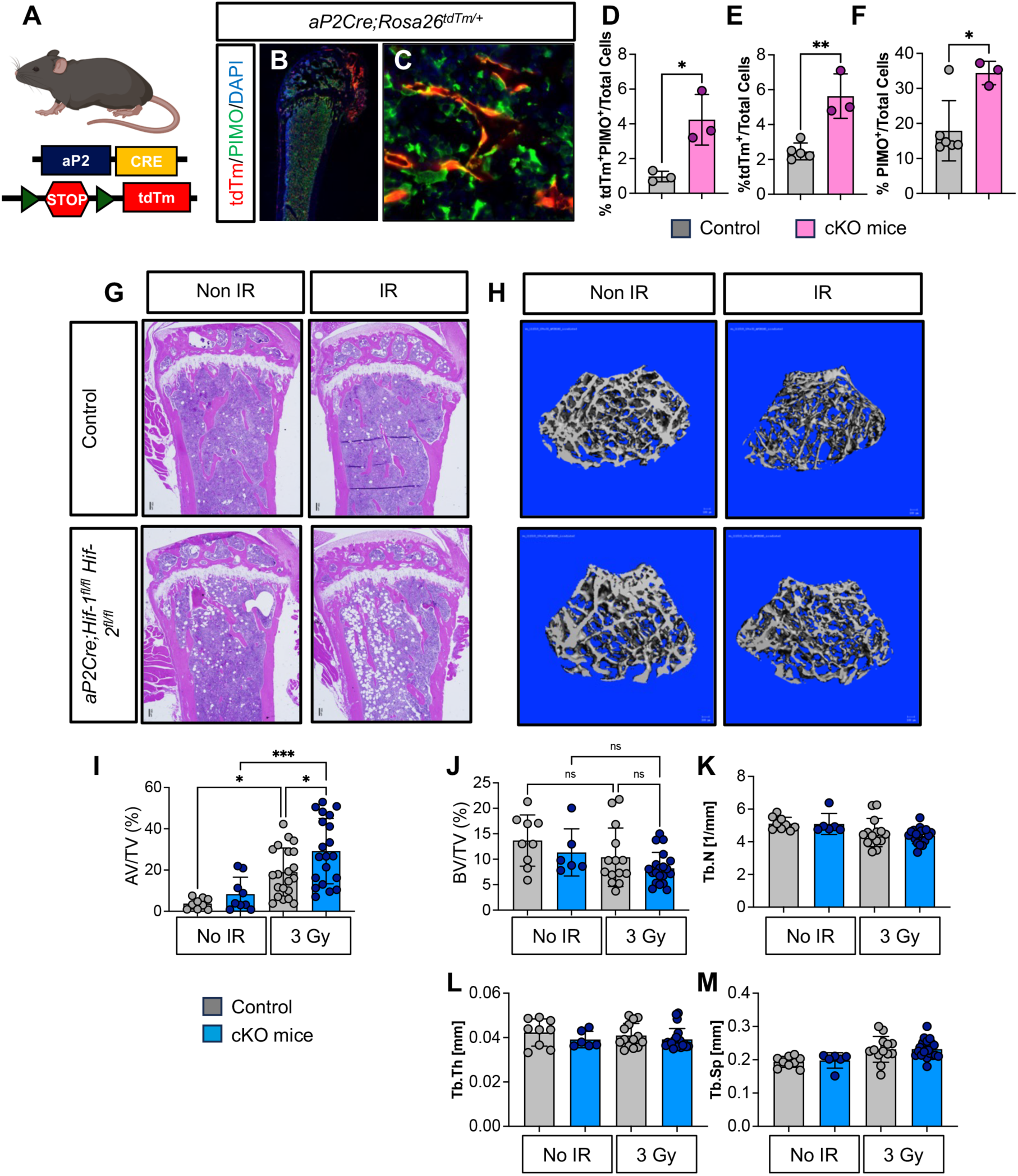
Conditional ablation of HIFα in aP2Cre expressing cells enhances radiation induced MAT. (**A**) Schematic representation of *aP2CreRosa26^tdTomato/+^*reporter lines. **(B-C)** Confocal images of long bones isolated from irradiated *aP2Cre;Rosa26^tdTomato/+^* mice stained for pimonidazole (PIMO) (green), and tdTomato (red). (**D-F**) Flow cytometry analysis of bone marrow flushes isolated from *aP2Cre;Rosa26^tdTomato/+^*mice treated with either 0 Gy or 3 Gy TBI. (**G**) Representative histological analysis and (**H**) microCT images of male *aP2Cre;Rosa26^tdTomato/+^*mice. microCT quantification of (**I**) osmium tetroxide staining and (**J)** bone volume/total volume (BV/TV) (**K**) trabecular number (Tb. N), (**L**) trabecular number (Tb.N.), (**M**) trabecular separation (Tb.Sp.) of bone volume and adiposity in *aP2Cre;Hif-1^fl/fl^;Hif-2^fl/fl^* cKO and *aP2Cre*(*-*)*;Hif-1^fl/+^;Hif-2^f/+^* control mice either in non-irradiated (non IR) or irradiated (IR) groups. Statistical analysis was performed using one-way ANOVA with Tukey’s post hoc test. Data are presented as means ± SEM. \**p* < 0.05, \*\**p* < 0.01, \*\*\**p* < 0.001, and \*\*\*\**p* < 0.0001.

### The majority of aP2Cre expressing cells do not label lipid laden adipocytes in MAT

Expression of the aP2Cre transgene has not been previously characterized in bone marrow adipose tissue which is unique in both function and developmental origins relative to other fat depots (48, 49). For this reason, we further characterized *aP2CreRosa26^tdTomato/+^*conditional reporter mice and evaluated the specificity of Cre mediated recombination of *aP2Cre* expressing cells in the hypoxic BME after radiation exposure. Intriguingly, despite robust expression of Tomato in the BME, demonstrating efficiency of aP2Cre expression, only 20% of Tomato^+^ cells were found to co-stain for perilipin after IR (**FIG 3A-D**). Confocal analysis revealed *aP2Cre* expressing cells were in close association but did not colocalize with, CD31^+^ and endomucin^+^ endothelial cells (**FIG 3G,H**). In support of these observations, flow cytometry analysis demonstrated that the majority of aP2Cre expressing bone marrow stromal cells did not express CD31, nor the macrophage marker CD11b **(FIG 3F)**. Collectively, our data suggests the majority of aP2Cre expressing cells in the BME represents a unique stromal cell population that are neither mature adipocytes, endothelial cells, nor macrophages.

**Figure 3.**
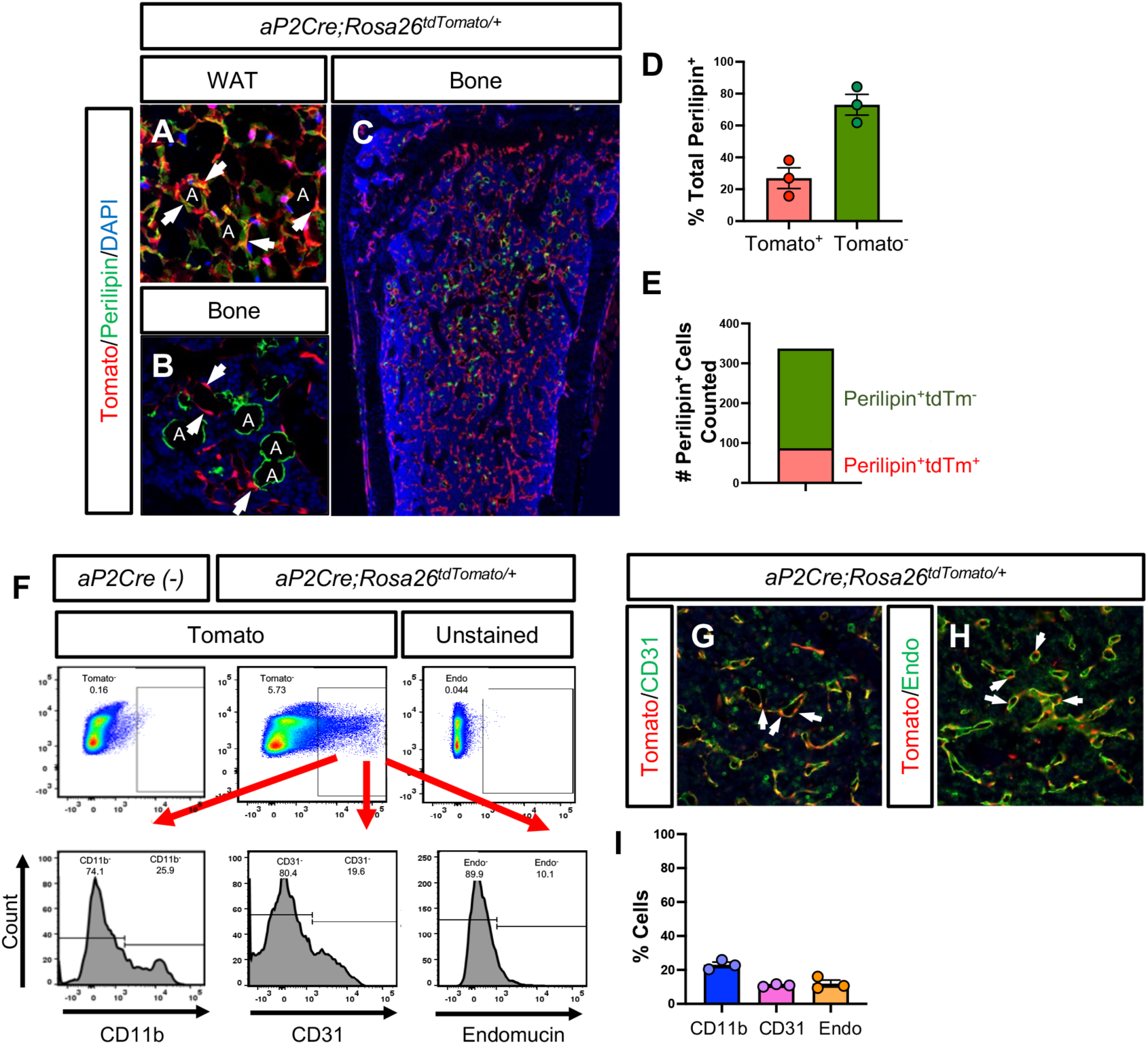
aP2Cre expressing cells do not co-stain with lipid laden adipocytes in the BME. Representative confocal images of **A)** white adipose tissue and **B)** marrow adipose tissue (**C**) and long bones isolated from IR *aP2Cre;Rosa26^tdTomato/+^*mice 7 days post IR stained for perilipin (green) and DAPI. A indicates adipocyte, white arrowheads indicate Tomato^+^ cells. **D-E)** Quantification of Tomato^+^perilipin^+^ cells in IR *aP2Cre;Rosa26^tdTomato/+^*mice 7 days post IR. **F**) Representative plots and quantification of flow cytometry analysis of cells isolated from long bones of *aP2Cre;Rosa26^tdTomato/+^*mice 7 days post IR and stained for CD11b, CD31, and endomucin **(G-H)** Confocal images of long bones isolated from irradiated *aP2Cre;Rosa26^tdTomato/+^* mice stained for **(G)** endomucin and (**H**) CD31. Statistical analysis was performed using one-way ANOVA with Tukey’s post hoc test. Data are presented as means ± SEM. \**p* < 0.05, \*\**p* < 0.01, \*\*\**p* < 0.001, and \*\*\*\**p* < 0.0001.

### aP2 Cre expressing cells label a unique population of stromal cells in the BME

Consistent with previous reports our preliminary data of *LepRCre;Rosa26^tdTomato/+^*and *AdipoQCre;Rosa26^tdTomato/+^* conditional reporter mice reveal Tomato^+^ cells within the bone marrow stoma in close association to the vasculature (**FIG 4A-C**) (50). Importantly, we show distinct expression patterns in the BME between the three murine models. Most notably, aP2Cre expressing cells are localized in the BM stroma along the periosteum and endosteal surfaces. In contrast, *AdipoQCre* and *LepRCre* expressing cells, while present in the stroma also label trabecular bone surfaces (**FIG 4A-C**). Moreover, confocal analysis of long bones isolated from *LepRCre;Rosa26^tdTomato/+^*, *AdipoQCre;Rosa26^tdTomato/+^* and *aP2Cre;Rosa26^tdTomato/+^* reporter lines and stained for perilipin^+^ adipocytes demonstrate that of unlike *aP2Cre expressing cells,* both *LepRCre* and *AdipoQCre* expressing cells directly contribute to bone marrow adipocytes after irradiation (**FIG 4D-L**) (25). Collectively, this suggests that aP2Cre labels a unique stromal cell population that is distinct from both *AdipoQCre and LepRCre expressing cells*.

**Figure 4.**
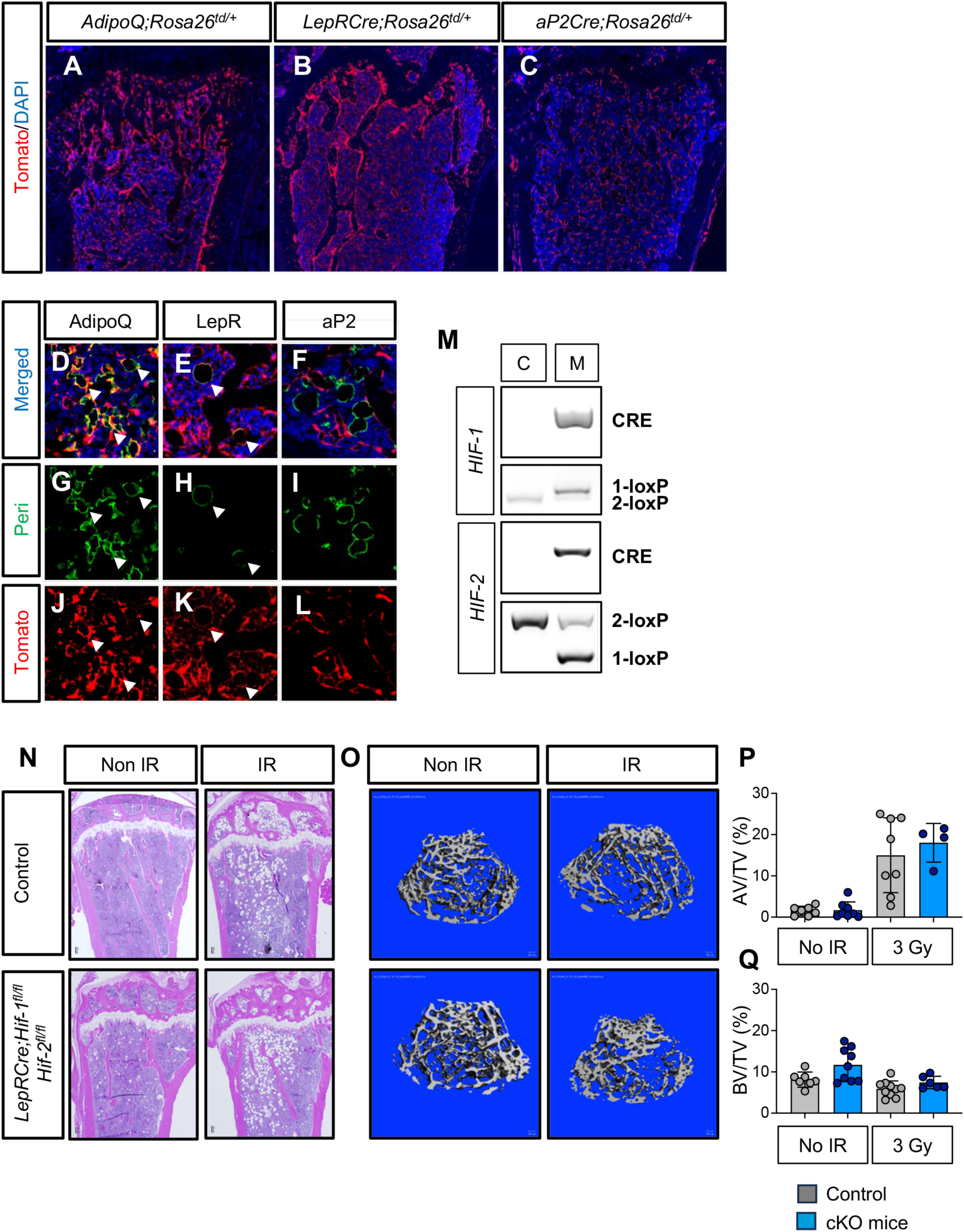
Characterization of bone marrow stromal cells. Confocal images of distal femurs isolated from 12-16 week old **(A)** *AdipoQCre;Rosa26^tdTomato/+^* **(B)** *LepRCre;Rosa26^tdTomato/+^* and **C)** *aP2Cre;Rosa26^tdTomato/+^*. Confocal images of distal femurs showing (**D-F)** merged images of **(G-H**) perilipin (green) and (**J-L**) tdTomato (red). DAPI (Blue). White arrowheads point to adipocytes. (**M**) qPCR analysis of genomic DNA isolated CFU-Fs isolated from *LepRCre*(*-*) (C) or *LepRCre;Hif-1^fl/fl^;Hif-2^fl/fl^* (M) mice. (**N**) Representative H&E and (**O**) microCT images and (**P**) microCT quantification of adipocyte volume/total volume and (**Q**) bone volume/total volume in *LepRCre;Hif-1^fl/fl^;Hif-2^fl/fl^* cKO or control mice either in non-irradiated (non IR) or irradiated (IR) groups. Statistical analysis was performed using one-way ANOVA with Tukey’s post hoc test. Data are presented as means ± SEM. \**p* < 0.05, \*\**p* < 0.01, \*\*\**p* < 0.001, and \*\*\*\**p* < 0.0001.

### Conditional ablation of Hif-1α and Hif 2α in LepRCre expressing cells does not MAT

For these reasons, to delineate the potential roles of HIF signaling in regulating radiation induced MAT within bone marrow stromal cells we generated *LepRCre;Hif-1^fl/+^;Hif-2^fl/+^* control and *LepRCre;Hif-1^fl/fl^;Hif-2^fl/fl^* conditional knock out mice and irradiated both groups. Importantly, as 94% of colony forming units fibroblastic (CFU-F) express LepR, efficient Cre-mediated recombination was confirmed by the presence of the 1Lox-P recombined allele from genomic DNA harvested from CFU-Fs isolated from *LepRCre;Hif-1^fl/fl^;Hif-2^fl/fl^* (**FIG 4M**) (21, 25). Intriguingly, in contrast to *aP2Cre;Hif-1^fl/fl^;Hif-2^fl/fl^* animals, histological analysis and microCT quantification of osmium tetraoxide staining revealed that of genetic ablation of HIFa transcription factors in LepRCre expressing cells did not attenuate the expansion of marrow adiposity in irradiated bones (**FIG 4N-Q)**. Collectively, this demonstrates that loss of HIF signaling in *LepRCre* expressing cells which directly contribute to MAT, does not impact adipocyte expansion after radiation exposure.

## DISCUSSION

Hypoxia/HIF signaling is essential for normal bone homeostasis and bone development with studies demonstrating that conditional ablation of HIF signaling in both chondrocytes, abd osteoblasts, resulting in limb defects and diminished bone mass. In contrast, the role of HIF signaling in bone marrow adipocytes has been largely unexplored. In these studies, we discovered that conditional ablation of HIFa in aP2Cre expressing cells resulted in an increase in radiation induced MAT expansion.

Here, we demonstrate that conditional ablation of HIF signaling in aP2Cre expressing cells does not affect MAT under homeostatic conditions but does increase MAT under radiation exposure. Based our labeling studies with *aP2CreRosa26^tdTomato/+^* reporter lines we show that the majority of aP2Cre expressing do not label lipid adipcyes, nor endotheial or macrophages. These findings suggest that in the bone marrow, aP2Cre cells mark a population of bone marrow stromal cells that may indirectly contribute to MAT expansion. Importantly, *in vitro* data of WAT show paracrine factors secreted by adipocyte regulatory cells in the stromal vascular fraction can influence both *de novo* adipogenesis and adipocyte hypertrophy (27). Intriguingly, many secreted factors that regulate adipogenesis such as *Agpat2, CaIII and Cd35* are putative HIF target genes (28, 29, 30, 31, 32, 33). Our preliminary animal studies highlight the need to profile and characterize unique stromal populations that indirectly support adipogenesis in the BME. Moreover, transcriptomic profiling of cells isolated from *aP2Cre;Hif-1^fl/fl^;Hif-2^fl/fl^* animals may also identify secreted factors that are HIF target genes which can be specifically targeted to prevent excessive MAT in irradiated bone.

Inhibition of the HIF signaling pathway in adipocyte regulatory cells in the BME enhance bone damage after exposure to radiation. These findings significantly contributes to the fields of bone biology, radiation biology, and the hypoxia/HIF signaling field by, 1) identifying a HIF-mediated mechanism that contributes to radiation damage to bone, and 2) characterizing bone marrow stromal cell populations that support adipocyte formation.

## MATERIALS AND METHODS

### Mice

All experimental procedures were approved by the Institutional Animal Care and Use Committee (IACUC) at Duke University. Mice were group-housed in animal care facilities under a 12 hr:12 hr light:dark cycle at 23±2°C with water and standard diet (LabDiet #5053, St. Louis MO) ad libitum. Mice were euthanized by CO_2_ asphyxiation for analyses. Experiments were performed on 12–16-week-old male and female mice on a C57Bl/6J background unless otherwise specified. *C57Bl/6J* (JAX:000664), *Rosa26tdTomato* (JAX:007909), *LepRCre* (JAX:008320), *Hif-2α^flox^* (JAX:008320), mouse strains were obtained from The Jackson Laboratory.

### Radiation treatment

Total-body irradiation was performed using an X-RAD 320 biological irradiator (Precision X-ray Inc., North Branford, CT) with a 2.5 mm aluminum, 0.1 mm copper filter and a field of view of 20 cm x 20 cm at a dose of 4 Gy. Dosimetry was measured with an ion chamber by the Radiation Safety Division at Duke University. For total-body irradiation, unanesthetized mice were placed in a ventilated cast acrylic pie cage (Braintree Scientific) without restraints and without shielding. The pie cage was centered on an acyclic slab to provide sufficient back scatter. All mice were placed 50 cm from the source and irradiated with 320 kVp, 12.5 mA X-rays.

Single-limb irradiation was delivered using a small animal irradiator, X-RAD 225Cx (Precision X-Ray Inc., North Branford, CT) or Small Animal Radiation Research Platform (SARRP, Xstrahl, Camberley UK). Prior to treatment, animals were anesthetized with nebulized isoflurane/O_2_ and positioned prone on the treatment stage. The irradiated limb received a single fraction of 8 Gy delivered AP/PA using a motorized collimator creating a 40 mm x 30-40 mm field at the treatment isocenter covering the entire right hindlimb, including the femur, tibia, and foot.

### RNA isolation and quantitative PCR (qPCR)

For BMSC cultures, total RNA was isolated and purified using the RNeasy Mini Kit (Qiagen, 74106) according to the manufacturer’s instructions. For bone homogenates, whole bone was extracted from animals, excess soft tissue was thoroughly removed, and samples were flash frozen in liquid nitrogen. Tissues were homogenized using a handheld mechanical homogenizer and total RNA was isolated and purified using the RNeasy Mini Kit. A total of 250ng of RNA was used to synthesized cDNA via reverse transcription reaction using the iScript gDNA Clear cDNA Synthesis Kit (Bio-Rad, 172-5035) according to manufacturer’s instructions.

Quantification of mRNA was determined by SYBR green chemistry (Bio-Rad, 172-5125) using the QuantStudio 6 Flex Real-Time PCR System (Applied Biosystems, 4484642). qPCR was performed in technical triplicates under the following conditions: denaturation at 95°C for 20s and 40 cycles of amplification at 95°C for 01s and 60°C for 20s. Gene expression was normalized to *Tbp* or *18s* mRNA and relative expression was calculated using a standard curve. Where indicated, gene expression was normalized to that of control samples. Primer sequences are listed in Table S1.

### Flow Cytometry

Flow cytometry analysis was conducted on bone marrow flushes obtained from 12–16-week-old control and experimental animals. Prior to staining, cells were filtered through a 70 μm nylon mesh. For live/dead cell exclusion, cells stained with the LIVE/DEAD Fixable Near-IR Dead Cell Stain Kit (Invitrogen, L10119) for 30 minutes at room temperature. For staining, antibodies were used at the following dilutions: Anti-mouse Endomucin eFluor 660 (1:100, Invitrogen, 50-5851-82), Affinity Purified Rabbit Anti-pimonidazole (1:100, Hypoxyprobe, HP3-1000Kit). A complete list of antibodies can be found in Table S2. All primary antibodies were incubated for 30 minutes at room temperature. Flow cytometry analysis was performed on a BD Fortessa X-20 (BD Biosciences) and analyzed using FlowJo software (version 10.6.1). Compensation was performed using OneComp eBeads Compensation Beads (Invitrogen, 01-1111-41) and ArC Amine Reactive Compensation Beads (Invitrogen, A10346) according to manufacturer’s instruction using appropriate fluorophore-conjugated antibodies. Living cells were gated for lack of anime reactive dye fluorescence with subsequent gating based on fluorescence-minus-one (FMO) and unstained controls.

### Histology

At indicated endpoints, bone tissues were harvested from mice, fixed in 10% neutral-buffered formalin for 72 hours and stored in 70% ethanol. Bone tissues were decalcified in 20% EDTA for 72 hours, processed using an automated tissue processor, and embedded in paraffin. Formalin-fixed paraffin embedded (FFPE) mid-sagittal tissue sections were taken at a 5 μm thickness and stained with hematoxylin and eosin (H&E) following standard protocols to visualize gross bone morphology. Adipocyte quantification was performed on H&E stained FFPE sections. For each biological sample, three 5 μm mid-sagittal sections were taken from the lateral, central, and medial regions of tibiae for quantification. For each section, total adipocyte number and average adipocyte size were quantified in a region of interest 1700 μm distal from the proximal growth plate. Marrow area was calculated as total area minus the bone surface.

### Micro-computed tomography (microCT) analysis

High resolution micro-computed tomography (VivaCT 80, Scanco Medical AG) was used for three-dimensional reconstruction and analyses of long bones for trabecular and cortical bone parameters. μCT scans for bone parameters were acquired using a 10 μm^3^ isotropic voxel size, 55 kVp peak X-ray tube potential, 200 ms integration time. A threshold of 338–1,000 and Gaussian filtering (σ=0.8, support=1) was used to segment mineralized bone from surrounding soft tissue. Analyses of trabecular bone included the measurement of 100 slices starting 50 slices below the secondary ossification center. Trabecular bone parameters assess include trabecular bone volume fraction (BV/TV, %), trabecular thickness (Tb.Th, mm), trabecular number (Tb.N, mm^−1^), and trabecular separation (Tb.Sp, mm). Cortical bone parameters were assessed by analyzing 50 slices in the mid-diaphysis at a set threshold of 300-1000 and Gaussian filtering (σ=0.8, support=1). Cortical bone parameters assessed include cortical area fraction (Ct.A/TA, %) and cortical thickness (Ct.Th, mm).

### Immunofluorescence

To obtain sections for immunofluorescent imaging, long bones were harvested from 12-16 week old animals. Harvested bones were fixed in 4% paraformaldehyde (PFA) for 24 hours followed by immersion with 30% sucrose for an additional 24 hours. Tissue samples were embedded in Cryomatrix (Thermo Scientific, 6769006) and cryosectioned at a 10 μm thickness using the CryoJane Tape transfer system. Primary antibody staining was performed with overnight incubation at 4°C using rat anti-Endomucin (Abcam, ab106100). Antibody binding was detected using appropriate secondaries: goat anti-rabbit IgG, Alexa Fluor 488 (1:200, Invitrogen, A-11008), rabbit anti-goat IgG, and Alexa Fluor 488 (1:200, Invitrogen, A-11078) with a 1 hr incubation at RT.

For visualization of pimonidazole, animals were injected with 60 mg/kg of pimonidazole HCl (Hypoxyprobe Omni Kit, HP3-1000Kit). Hypoxic protein adducts were detected with overnight incubation of rabbit antisera (supplied in Hypoxyprobe kit) at 4°C and detected with Alexa Fluor 488-conjugated goat anti-rabbit secondary antibody (1:200, Invitrogen, A-11008) 1 hr incubation at RT.

Fluorescent images were captured by a Zeiss LSM 780 inverted confocal microscope interfaced with the Zen Black software (Carl Zeiss Microimaging LLC, Thornwood, NY)

## Author contributions

Conceptualization: WG, CW

Methodology: CAJ, WG, CW

Investigation: CAJ, WG, KAG, CP, AS

Visualization: CAJ, WG,

Funding acquisition: CW

Project administration: CW

Supervision: CW

Writing – original draft: CAJ, CW

Writing – review & editing: CAJ, CW

## Data and materials availability

All data are available in the main text or the supplementary materials.

