## Supplementary Figures for "Hypoxia/HIF Signaling Negatively Regulates Bone Marrow Adiposity after Radiation Exposure"

### SUPPLEMENTARY FIGURE 1

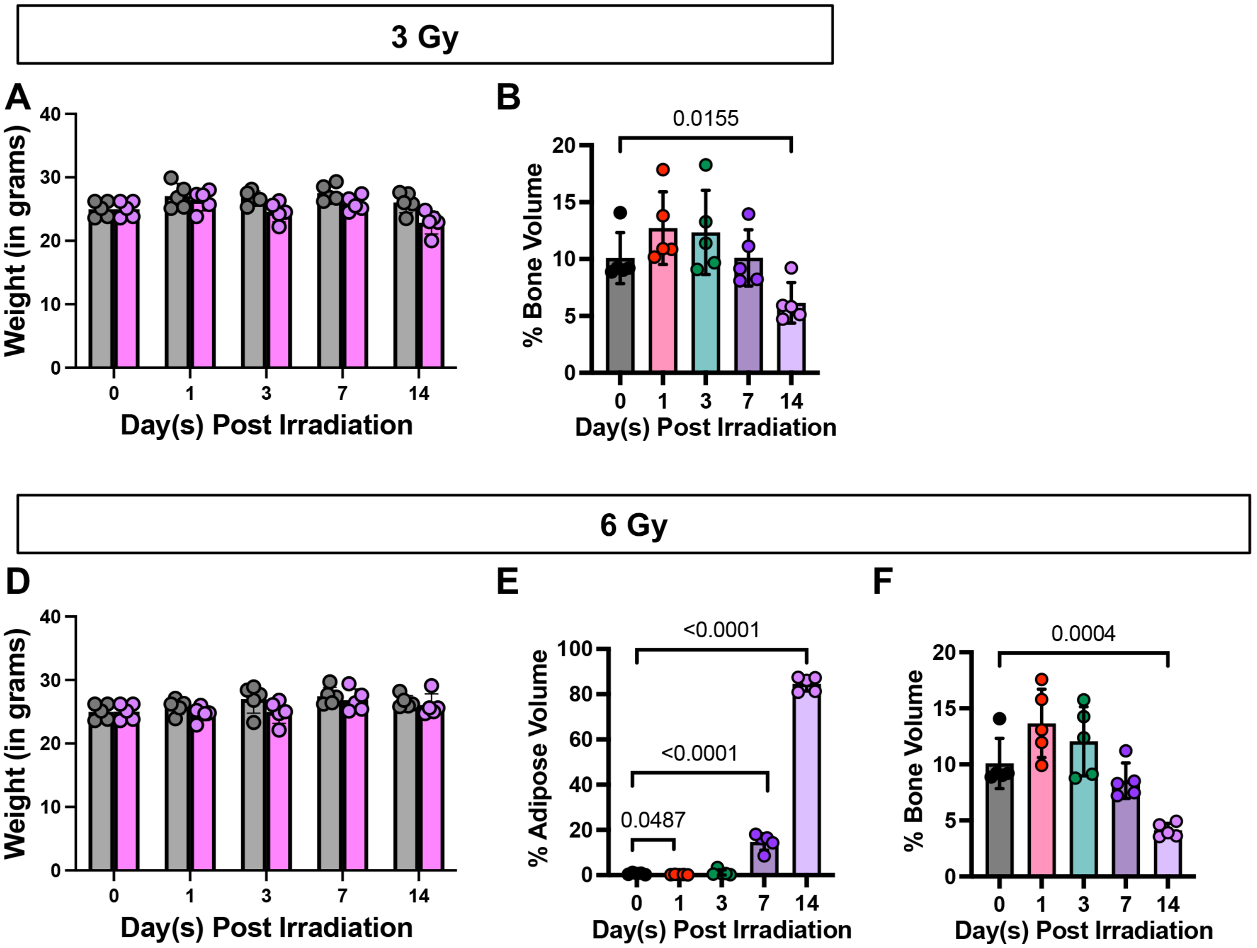

**Supplementary Figure 1** (**A**) Weights of mice and (**B**) microCT quantification of bone volume of proximal tibias and 1, 3, 7, and 14 days post 3 Gy TBI. (**C**) Weights of mice (**D**) microCT quantification of bone volume of proximal tibias and (**E**) osmium tetroxide staining of distal femurs 1, 3, 7, and 14 days post 6 Gy TBI.

### SUPPLEMENTARY FIGURE 2

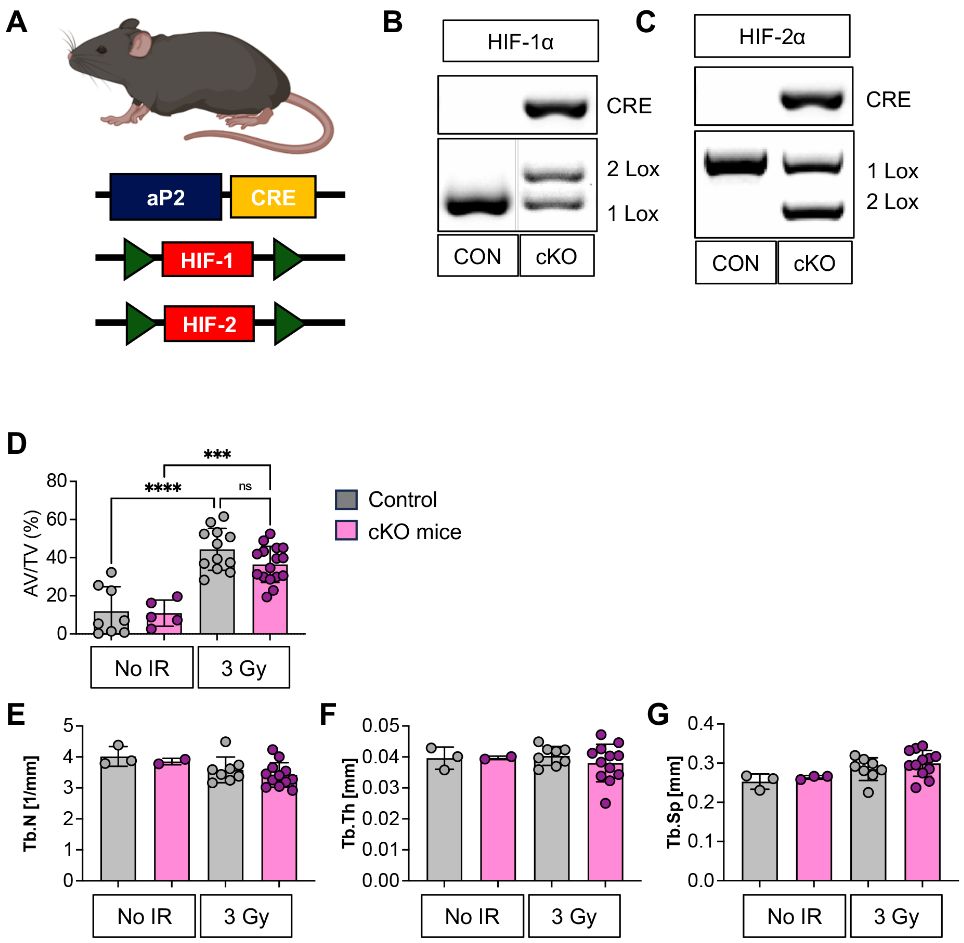

**Supplementary Figure 2**

(**A**) Schematic representation of *aP2Cre;Hif-1$^{fl/fl}$;Hif-2$^{fl/fl}$* cKO reporter lines. (**B,C**) qPCR analysis of genomic DNA isolated from adipocytes isolated from *aP2Cre(-)* control (C) or *aP2Cre;Hif-1$^{fl/fl}$;Hif-2$^{fl/fl}$* cKO (M) mice. (**D**) microCT quantification of osmium tetroxide staining of femurs isolated from female *aP2Cre(-)* control (C) or *aP2Cre;Hif-1$^{fl/fl}$;Hif-2$^{fl/fl}$* cKO mice either in non-irradiated (non IR) or irradiated (IR) groups (**E-G)** microCT quantification of proximal tibias isolated from female *aP2Cre(-)* control (C) or *aP2Cre;Hif-1$^{fl/fl}$;Hif-2$^{fl/fl}$* cKO mice either in non-irradiated (non IR) or irradiated (IR) groups**.** (**E**) trabecular number (Tb. N), (**F**) trabecular thickness (Tb.N.), and (**M**) trabecular separation (Tb.Sp.) control mice. Statistical analysis was performed using one-way ANOVA with Tukey's post hoc test. Data are presented as means ± SEM. *$p < 0.05$, **$p < 0.01$, ***$p < 0.001$, and ****$p < 0.0001$.
